# Elicitor specific mechanisms of defence priming in oak seedlings against powdery mildew

**DOI:** 10.1101/2024.09.20.613659

**Authors:** Rosa Sanchez-Lucas, Jack L. Bosanquet, James Henderson, Marco Catoni, Victoria Pastor, Estrella Luna

## Abstract

Defence priming sensitises plant defences to enable a faster and/or stronger response to subsequent stress. Various chemicals can trigger priming; however, the response remains unexplored in oak. Here, we characterise salicylic acid (SA)-, jasmonic acid (JA)-, and β-aminobutyric acid (BABA)-induced priming of oak seedlings against the causal agent of powdery mildew (*Erysiphe alphitoides,* PM). Whilst JA had no effects, BABA and SA enhanced resistance by priming callose deposition and SA-dependent gene expression, respectively. Untargeted transcriptome and metabolome analyses revealed genes and metabolites uniquely primed by BABA, SA, and JA. Enrichment analyses demonstrated a limited number of pathways differentiating the three treatments or the resistance-inducing elicitors BABA and SA. However, a similar mode of action between BABA and JA was identified. Moreover, our analyses revealed a lack of crosstalk between SA and JA. Interestingly, priming by BABA was linked to alkaloid, lignan, phenylpropanoid, and indolitic compounds biosynthesis. Moreover, integration of the omics analyses revealed the role of ubiquitination and protein degradation in priming by BABA. Our results confirm the existence of chemical-induced priming in oak and has identified specific molecular markers associated with well-characterised elicitor.

## Introduction

Forests cover 31% of the Earth’s land surface (FAO, 2022) and have an estimated global value of US$150 trillion (WWF, 2023). For the UK alone, forests directly generate £4.9 billion each year (DEFRA, 2018). Forests also have major social, cultural, and religious value, for example as historic landmarks that link generations (Blicharska and Mikusinski, 2014). Moreover, forests support much of the Earth’s biodiversity (∼80% of terrestrial diversity (Aerts and Honnay, 2011)) and act as a major store of carbon (Bonan, 2008). Therefore, efforts to limit deforestation, manage existing forests, and restore or re-establish those that have been lost, are of critical importance (Griscom et al., 2017; FAO, 2022; Kauppi et al., 2022).

A key tree genus in the northern hemisphere, *Quercus* dominates much of the world’s forests (Mölder et al., 2019). The UK has two native species of oak, *Quercus petraea* and *Quercus robur* (Mitchell et al., 2019; Mölder et al., 2019), which make up 16% of the countries’ broadleaf woodland and have an estimated annual value of £320 million (DEFRA, 2018). Unfortunately, oak trees remain threatened by various stresses that have led to their deteriorating health and survival over the past 100 years (DEFRA, 2018). Of these threats, oak powdery mildew (PM) is particularly problematic in seedlings (Mougou et al., 2008). The primary causal agent is *Erysiphe alphitoides*, to which *Q. robur* is more susceptible than *Q. petraea* (Marçais and Desprez-Loustau, 2014). Young or heavily infected leaves suffer necrosis and deformation which can cause decreased growth, reduced photosynthesis, and death (Hajji et al., 2009; Marçais and Desprez-Loustau, 2014; Sanchez-Lucas et al., 2023). Ultimately, this reduces the ability of young trees to compete, making the disease a major bottleneck for oak woodland regeneration (Demeter et al., 2021). Control of PM in tree nurseries currently relies heavily on chemical fungicides, which is limited due to environmental toxicity (Nayana and Ritu, 2017). Thus, a better understanding of the oak immune system is needed for the development of sustainable strategies that reduce the impact of PM.

The plant innate immune system consists of two interconnected layers. Upon initial infection, cell-surface pattern recognition receptors (PRRs) detect pathogens and activate pattern-triggered immunity (PTI). The second layer involves intracellular nucleotide-binding domain leucine-rich repeat receptors (NLRs) that recognise pathogen virulence molecules (effectors) and trigger effector triggered immunity (ETI) (Dodds and Rathjen, 2010; Yuan et al., 2021b; Yu et al., 2024). PTI provides broad-spectrum resistance against non-adapted pathogens (i.e. non-host resistance) thanks to various defence responses including, production of antimicrobial compounds, reactive oxygen species (ROS) burst, and callose deposition (Dodds and Rathjen, 2010; Yu et al., 2017). ETI involves many similar responses to PTI, but the main distinguishing feature is a form of localised programmed cell death known as the hypersensitive response (Coll et al., 2011;Yuan et al., 2021b). PTI and ETI are tightly linked and work synergistically to activate defence responses and promote resistance (Ngou et al., 2021; Yuan et al., 2021a; Yu et al., 2024). Regulation of, and crosstalk between, defence responses is facilitated by, among other mechanisms, phytohormones such as salicylic acid (SA) and jasmonic acid (JA) (Yu et al., 2024). JA, in conjunction with ethylene (ET), is often associated with responses to necrotrophic pathogens, while SA is commonly linked to immunity against biotrophs and hemi-biotrophs (Glazebrook, 2005; Shigenaga et al., 2017). In many herbaceous plants, an antagonism between SA and JA has been reported: activation of SA-dependent defences suppresses JA-dependent defences and *vice versa* (Pieterse et al., 2012; Shigenaga et al., 2017; Yu et al., 2024).

Plants also possess the capacity to acquire immunological memory in the form of priming of defence (Martinez-Medina et al., 2016). Following exposure to a priming stimulus, the sensitivity of the plant immune system increases, which facilitates a faster and stronger induction of defence responses upon subsequent stress (Mauch-Mani et al., 2017). Biological mechanisms behind priming include changes in the levels of signalling molecules or metabolites and epigenetic changes that alter gene expression (Cooper and Ton, 2022; Hannan Parker et al., 2022). The primed state can last throughout the plant life cycle and may also be passed between generations via epigenetic inheritance (Mauch-Mani et al., 2017).

The best characterised priming response is systemic acquired resistance (SAR), which gives long-lasting, broad-spectrum resistance and relies on SA signalling. SAR is thought to be triggered by HR during infection and is often associated with pathogenesis-related (PR) gene expression (Conrath, 2006). Beneficial soil microbes can also trigger priming via induced systemic resistance (ISR), or mycorrhiza induced resistance (MIR). ISR is primarily associated with JA- and ET-mediated defences whilst MIR has been linked to both JA and SA (Cameron et al., 2013; Benjamin et al., 2022). JA-dependent defences and volatile organic compounds (VOCs) are also linked to a form of priming called herbivore-induced resistance (HIR) (Pieterse et al., 2014; Erb et al., 2015). Priming relies on complex chemical signalling cascades, which can be manipulated by treating plants with synthetic or natural chemicals to establish priming of defence. For example, exogenous SA treatment mimics SAR (Bawa et al., 2019), whilst JA treatment can mimic ISR or HIR (Arévalo-Marín et al., 2021; Bhavanam and Stout, 2021). Another well characterised priming chemical is β-aminobutyric acid (BABA), which triggers a process known as BABA-induced resistance (BABA-IR) that primes broad-spectrum resistance to various plant pathogens (Cohen et al., 2016; Wilkinson et al., 2018), including PM in oak (Sanchez-Lucas et al., 2023). BABA-IR can overlap with SA- and/or JA-/ET-mediated defences or may occur independently of either (Zimmerli et al., 2000; Ton and Mauch-Mani, 2004; Jakab et al., 2005). In Arabidopsis, binding of BABA to the aspartyl-tRNA synthetase IBI1 promotes the defence responses associated with BABA-IR (Luna et al., 2014). For example, IBI1 activates VOZ1/2 transcription factors, which induce abscisic acid (ABA)-dependent callose deposition to slow pathogen invasion and spread (Schwarzenbacher et al., 2020). A limitation of BABA is that it can impair plant growth at concentrations required for BABA-IR in certain species (van Hulten et al., 2006; Wu et al., 2010; Luna et al., 2016). However, this appears not to be the case in oak seedlings (Sanchez-Lucas et al., 2023), suggesting that BABA could control PM in tree nurseries without negatively impacting seedling growth.

Most research into priming has been conducted in short-lived model plants and has focused on elucidating conserved molecular mechanisms. Whilst trees share certain features with all plants, major differences, including their larger size, longer lifespans, and wood production, mean it is likely that priming mechanisms can differ in these species (Mageroy et al., 2020a). Of the few studies that have examined priming phenomena in trees, abiotic, rather than biotic, stresses have often been the focus (Amaral et al., 2020). However, several studies have demonstrated priming against biotic stress in trees (Bittner et al., 2019; Camisón et al., 2019; Mageroy et al., 2020a; Mageroy et al., 2020b; Martínez-Arias et al., 2021). Moreover, whilst research remains limited, induced resistance has been shown to occur in oak species. For example, beneficial *Streptomyces* rhizobacterium enhance resistance to PM (*Microsphaera alphitoides*) in *Q. robur* (Kurth et al., 2014). However, whether the induced resistance observed is mediated by priming remains unknown.

Omics allows for high-throughput, high-resolution, unbiased, characterisation of complex biological systems (Kan et al., 2017), making them invaluable for understanding the complex mechanisms behind priming of defence. When combined, omics (e.g. transcriptomics, metabolomics, proteomics) give holistic insight into the ‘prime-ome’ (Balmer et al., 2015). Omics have already advanced our understanding of both SAR and ISR in model plants (Razzaq et al., 2023), where they are especially useful for putative identification of novel priming molecules which can be confirmed biochemically. Omics have also been used to better understand induced resistance in non-model species like trees. For example, RNA-seq has shown that ISR in oak shares some similarities with Arabidopsis (upregulation of phenylpropanoid biosynthesis genes) but also some differences (the involvement of both JA/ET and SA pathways) (Kurth et al., 2014). However, large-scale omics experiments to unravel priming mechanisms in oaks are lacking. Priming presents a novel way to protect plants against fungal diseases, however our understanding is much better in herbaceous plants than in forest trees. This study aims to characterise chemical-induced priming during infection of the most susceptible UK oak species, *Q. robur*, with PM. Little is known about the genes and metabolites that may be involved, thus untargeted transcriptomics and metabolomics analyses were performed to putatively identify novel biomarkers of the priming response in oak which may enhance protection of seedlings against this limiting pathogen.

## Material and Methods

### Plant material and growth conditions

Acorns of *Quercus robur* UK provenance 405 (Hubert and Cundall, 2006) were sourced from the tree seed company Forestart (https://www.forestart.co.uk) and were germinated according to existing protocols (Simova-Stoilova et al., 2015; Simova-Stoilova et al., 2018; Sanchez-Lucas et al., 2023). After 72h, germinated acorns were transferred into individual root trainers (Maxi Rootrainers, Haxnicks, RT230101) containing 400 mL of Scott’s Levington M3 Advance Pot & Bedding soil. A small part of the experiments were grown with Levington Advanced F2 soil (peat free). Germinated acorns were then grown in a glasshouse compartment at 16/8h light day/night (∼200 μmol photons m-2s-1), 20_/18_ cycle, 42% HR and irrigated to field capacity throughout the experiment.

### Elicitor treatment

Three elicitors were used: salicylic acid (SA), jasmonic acid (JA) and β-aminobutyric acid (BABA). Treatments were performed on 3-month-old seedlings. BABA was soil-drenched to a final soil concentration of 5mM as previously described (Sanchez-Lucas et al., 2023). SA and JA were sprayed at concentrations of 5mM and 500μM, respectively. Water was used as control in both soil drench and spray treatments, along with being applied to each elicitor to equalise the amount of water in the soil or on the leaves. All spraying solutions contained 0.002% ethanol and 0.05% Silwet to allow full leaf coverage.

### Pathogen infection and disease scoring

Plants were inoculated with PM causal agent, *Erysiphe alphitoides*, 7 days post elicitor treatment, by spraying leaves with 1.5×10^6^ spores/mL. Mock-inoculated plants were treated with water. All inoculations contained 0.05% of Silwet to allow full leaf coverage. Post-inoculation, all plants were grown under high humidity by covering the plants with plastic bags.

Disease scoring was performed at 14- and 30-days post infection (dpi). Between 9 and 12 plants were scored per elicitor. At 14 dpi the mean number of individual colonies across 10 leaves was recorded for each treatment. At 30 dpi, due to increased fungal growth, individual colonies were no longer visible. At this point, the number of leaves that fell into four different categories of fungal colonisation were quantified: 1) healthy; 2) <50% mycelium leaf coverage; 3) >50% mycelium leaf coverage; 4) necrosis and tissue collapse. A Disease Severity Index (DSI) based on the 4 categories was calculated using the formula:

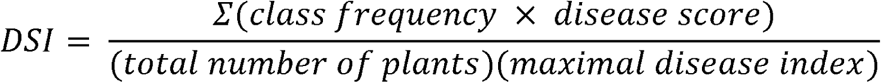

For field conditions, one month-old seedlings were grown and treated with elicitors as described above. Seven days post elicitor treatment plants were transported to the Birmingham Institute of Forest Research (BIFoR) Free Air CO2 Enrichment (FACE) facilities in July 2020, where plants grew in ambient Array 3 (Hart et al., 2020) and placed into plastic net cages until the end of the experiment. Spores from strains local to the facilities were collected in water from young oak trees an hour before inoculation (which was performed as described above). In field conditions, disease scoring was only possible at 30 dpi, with DSI measured as above.

### Plant growth parameters

Relative growth rates (RGR) were calculated for height, main stem diameter, and leaf length for mock treatments between the timepoints 7 days before elicitor treatment (t1) and 21 days post treatment (t2). RGR was calculated using the formula:

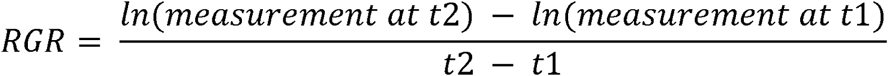

The number of leaves per plant at t2 was used to further characterise plant growth upon elicitor treatment.

### Fluorescence microscopy quantification of callose deposition

Leaf-discs (25mm^2^) from mock- and PM-inoculated plants (7 replicates per treatment) at 0 and 1 dpi were subjected to double staining with aniline blue and calcofluor to visualise callose and pathogen mycelia, respectively, as modified from Luna et al., 2011. Briefly, leaf-discs were bleached in 96% ethanol, before discarding the ethanol and incubating them in 0.07M phosphate buffer (pH 9) overnight to allow rehydration. Afterwards, aniline blue (Acros Organics, CAS 28983-56-4) and calcofluor (Fluorescent Brightener 28, Sigma, F3543) staining was performed by incubating leaf-discs in the dark with 0.05% aniline blue and 0.025% calcofluor at a ratio of 3:1 for 15 minutes. Then, upon removal of the staining solution, leaf-discs were incubated in the dark with 0.05% aniline blue overnight. Finally, slides were prepared in a matrix of 0.05% aniline blue.

Fluorescence microscopy was performed with the 10X magnification lens of a GX Microscope L Series (GT Vision Ltd) coupled to a GX camera. Images were recorded with GX Capture 8.5 (GT Vision Ltd) software at an image resolution of 96 dpi (dots per inch). Total amount of callose and callose-induced by pathogen was quantified using Photoshop version 22.2.2. For the assessment of callose deposition-induced by the elicitors, measurements of the number of pixels corresponding to callose deposition were made. The mean area of callose deposition per leaf was then calculated. Effective callose induced by the pathogen was assessed by categorising areas of pathogen growth into different categories of callose deposition: I - pathogen presence/no callose; II- pathogen presence/ineffective callose; III- pathogen presence/effective callose; IV- pathogen spore present/effective callose. An index was calculated following the formula for the DSI described above.

### Statistical analyses

Boxplots and statistical testing were done in R v4.3.1 (R_Core_Team, 2023). Boxplots were drawn using the ggplot2 package (Wickham, 2016). Data were first examined to confirm they fitted the assumptions for ANOVA using the stats (R_Core_Team, 2023) and olsrr packages (Hebbali, 2024). Normality of the residuals was checked with the Shapiro-Wilk test (p > 0.05). Homogeneity of variances was checked with a Levene’s test within the CAR package (Fox and Weisberg, 2019). For non-normal data a Kruskal-Wallis test was performed, followed by Dunn’s post hoc test using the FSA package (Ogle et al., 2023). For data that met the normality and homoscedasticity assumptions a one-way ANOVA was performed, followed by a Tukey post hoc test using the Multcomp package (Hothorn et al., 2008). For data that met the normality but not the homoscedasticity assumptions a Welch ANOVA was performed, followed by a Dunnett’s T3 post hoc test using the PMCMRplus package (Pohlert, 2023).

### Sample collection for omics analyses

Parallelly, samples for both omics analyses were collected at 0, 1, and 2dpi, by snap-freezing mock and infected leaves in liquid nitrogen and stored at -80°C until processing. Material was collected from 4 biological replicates (seedlings) per group giving a total of 80 samples for each omics analysis.

### Transcriptomic analysis

#### RNA extraction, library preparation and mRNA sequencing

Total RNA was extracted using the Macherey-Nagel, Mini-Plant RNA extraction kit following manufacturer instructions with modifications for leaf tissue. Briefly, a 1:1 combination of PL1 and PL2 was applied during the lysis step. Centrifuge times were duplicated during the extraction and two additional washing steps were incorporated. RNA quality was tested with nanodrop and gel electrophoresis.

mRNA sequencing was conducted on RNA extracted from all the conditions. Library preparation and sequencing was conducted by Novogene using NovaSeq 6000 PE150. Around 40 million paired reads were generated per sample. An average of 93.63% of nucleotides per sample had a Phred quality score of > 30. The quality of samples was assessed using FAST QC (0.11.5-Java-1.8.0_74) (Andrews, 2019). Adapters were removed from samples using Trimmomatic (version 0.39) (Bolger et al., 2014). Reads were aligned to the British oak genome (https://www.oakgenome.fr/) using Hisat2 (version 2.2.1) (Kim et al., 2019). Samtools (version 1.12) (Danecek et al., 2021) was used to sort and index Sam files into Bam format. Read counts were generated using HTSeq (version 0.13.5) (Putri et al., 2022) using default thresholds.

#### Global data visualisation

Global data visualisation was performed in R v4.3.1 (R_Core_Team, 2023). As recommended in the DESeq2 (Love et al., 2014) documentation a regularised log (rlog) transformation was performed on the raw counts prior to visualisation (see Love et al., 2024). Using the package mixOmics v6.1.2 (Rohart et al., 2017) principal component analysis (PCA) and Partial Least Squares Discriminant Analysis (PLS-DA) were performed to visualise differences between the various treatments.

#### Analysis of differential gene expression

Analysis of differential gene expression was performed in R v4.3.1 (R_Core_Team, 2023). DESeq2 v1.42.1 package was used to analyse the raw counts data without rlog transformed (Love et al., 2024). The DESeq algorithm estimates size factors and dispersion and performs the Wald significance test following negative binomial GLM fitting to find differentially expressed genes.

An adjusted Wald test p-value of 0.05 was used to select significant differentially expressed genes for both mock- and PM-inoculated treatments. Data was mean-centred and divided by the standard deviation before using Dendextend v1.17.1 (Galili, 2015) and ComplexHeatmap v2.18.0 (Gu et al., 2016; Gu, 2022) to plot dendrograms and heatmaps visualising the clustering of significant DEGs. For the isolation of primed genes at 1dpi and 2dpi we compared each elicitor + mock treatment against elicitor + PM treatment and selected those differentially expressed genes solely associated with PM infection. These were subsetted into up- and downregulated genes based on a log2 fold-change of >1 or <-1, respectively. Venn diagrams were then drawn using VennDiagram v1.7.3 (Chen and Boutros, 2011) to compare the effect of the different elicitors on priming of gene expression.

#### GO term annotation and enrichment analysis

GO term annotation was performed on the oak genome assembly PM1N gene prediction (.gff) file (https://www.oakgenome.fr/). Transdecoder v5.7.1 (Haas, 2023) was used to extract long ORFs. BLAST+ v2.14.0 (BLASTp homology) (Camacho et al., 2009) and HMMER3 v3.3.2 (protein domain homology) (Eddy, 2011) were used to check for homology and refine the ORF search. The final GO annotations were produced using Trinotate v4.0.2 (Bryant et al., 2017), which was run with default options using DIAMOND v2.1.8 (Buchfink et al., 2021) for efficient BLAST searching. In total, 21,710 transcripts were mapped to GO terms.

GO enrichment analysis was performed for the identified primed genes for all of the elicitors in R v4.3.1 (R_Core_Team, 2023) using topGO v2.54.0 (Alexa and Rahnenfuhrer, 2023). The Weight01 algorithm and Fisher’s exact test were used to determine significant enrichment of GO terms. Summary tables for the data were produced by selecting the top 10 most significant (p≤0.05) GO terms with at least 10 gene hits.

### RT-qPCR quantification of PR1 expression

RNA samples were subaliquoted for RT-qPCR to determine changes in *PATHOGENESIS RELATED PROTEIN 1* (*PR1*) following elicitor treatment and PM infection at 1dpi. cDNA synthesis was performed using SuperScript IV VILO Master Mix (ThermoFisher, Catalogue reference 11756050) following manufacturer instructions. PowerUp™ SYBR™ Green Master Mix (Applied Biosystems™, Catalog number: A25741) was used following manufacturer instructions. QuantStudio™ 5 Real-Time PCR System, 384-well (Applied Biosystems™, Catalog number: A28140) was run for 40 cycles of qPCR. Ct values were analysed using the 2^-ΔΔCT^ method (Livak and Schmittgen, 2001). Using His3.2 and EF1 as reference genes, average fold change in PR1 expression was calculated. Primers were designed based on a NCBI nucleotide database (https://www.ncbi.nlm.nih.gov/nucleotide/) search using *PR1* and *Quercus* as filters. Primer3web v4.1.0 (Kõressaar et al., 2018) was employed to design primer pairs of 17-20 nucleotides, 40-60% GC content, and T_m_ of 57-63°C. OligoCalc (Kibbe, 2007) was used to evaluate for potential secondary structures. Primer specificity was checked using Primer-BLAST (Ye et al., 2012). PR1 primer sequences were: PR1-Fw: CGCTGTGAACATGTGGGTAG and PR1-Rv: TGTTGCATCGAACTTTGGCA. Primers used for the two internal reference genes were His3.2-Fw GCTCTTCGAGGACACCAATC and His3.2-Rv TAAGCCCTCTCGCCTCTGAT; EF1-Fw TTGTGCCGTC CTCATTATTGACT and EF1-Rv TCACGGGTCT GACCATCCTT.

### Metabolomic analysis

#### Metabolite extraction and LC-QTOF analysis

Plant material was lyophilised before extraction. Metabolite extraction was performed using the methanol extraction protocol described in Pastor et al., 2018. Briefly, after the addition of 30% methanol, samples were incubated on ice for 30 minutes prior to centrifuge. The supernatant was filtered (using 0.2µm regenerated cellulose, Teknokroma) prior to LC-MS/MS.

Untargeted metabolomics was performed using the Acquity UPLC system (Waters, Milford, MA, USA) interfaced to a hybrid quadrupole time-of-flight mass spectrometer (Q-TOF MS Premier). LC was performed with a C-18 column (Kinetex C18 analytical column, 1.7 µm particle size, 50 mm□×□2.1 mm (Phenomenex)).

#### Data processing

Raw LC-MS/MS spectra files were extracted using Masslynx 4.2 as previously described (Colombo et al., 2004; Manresa-Grao et al., 2024) before being filtered in R v4.3.1 (R_Core_Team, 2023) using the XCMS package with the parameters defined on the Centwave (Smith et al., 2006; Tautenhahn et al., 2008). Mass traces with at least 4 peaks with an intensity ≥200 and width of 5-12 seconds were retained. MS1 peak intensity values were normalised by dry weight per sample before further analysis. Adduct and isotope correction, followed by merging the positive and negative ionisation mode data, was performed using MarVis-Filter within the MarVis-Suite software (Kaever et al., 2015).

#### Global data visualisation and statistical analysis

Global data visualisation was performed using Metaboanalyst 6.0 (Pang et al., 2021). Principal Component Analysis (PCA) and Partial Least Squares Discriminant Analysis (PLS-DA) were used to group putative metabolites under different treatments. A Kruskal-Wallis was then used to determine metabolites that varied significantly (p ≤ 0.01) between treatments. This was performed without false discovery rate (FDR) correction. Using R v4.3.1 (R_Core_Team, 2023), data was mean-centred and divided by the standard deviation. The Dendextend v1.17.1 (Galili, 2015) and ComplexHeatmap v2.18.0 (Gu et al., 2016; Gu, 2022) packages were then used to plot dendrograms and heatmaps visualising the clustering of significant metabolites (p ≤ 0.01).

#### Selection of primed metabolites

Mean fold-change in metabolite abundance (across the 4 replicates) was calculated for each treatment in the merged dataset. Those metabolites with a p-value ≤0.05 and fold-change of ≥2 or ≤-2 were used in subsequent analysis. For 1 and 2 dpi, primed metabolites were isolated following the previous approach by comparing elicitor + mock and elicitor + PM infected samples and selecting those significant metabolites solely associated with infection. Venn diagrams were drawn using the VennDiagram v1.7.3 (Chen and Boutros, 2011) package in R v4.3.1 (R_Core_Team, 2023) to compare the effect of the different elicitors on metabolite priming.

#### Pathway enrichment analysis

Primed metabolites underwent pathway analysis using the MarVis-Suite software (Kaever et al., 2015). Data was clustered using MarVis-Cluster and subsequent pathway analysis was performed with MarVis-Pathway without clustering selection. Data was ranked and matched to several metabolite libraries (the publicly available KEGG database for *Populus trichocarpa* and internal libraries kindly provided by Dr Pastor’s group (Manresa-Grao et al., 2022; Manresa-Grao et al., 2024)). Thresholds for differences in m/z and RT were applied, 0.01 Dalton and 5 seconds, respectively. Entry based enrichment analysis calculated p-values based on a hypergeometric distribution, which were then adjusted using FDR (Benjamini-Hochberg) correction. Pathways with a p-value of ≤0.05 were deemed significantly enriched.

### Hormone levels measurement

Exact neutral masses for plant hormones and their conjugates were isolated from the obtained mass spectra using the MassLynx V4.2 Software and as described before (Colombo et al., 2004; Manresa-Grao et al., 2024). This was done for the monoisotopic masses of salicylic acid (SA, 138.031693 Da;MS2: 65.0384/ 93.033/ 94.0358/137.0231), jasmonic acid (JA, 210.125595 Da;MS2: 165.129/209.0727/209.1175), jasmonic acid-valine (JA-Val, 309.194008 Da;MS2: 165.129/209.0727/209.1175/118.0868) and jasmonic acid-isoleucine (JA-Ile, 323.20966 Da;MS2: 165.129/209.0727/209.1175/44.052/45.035/53.042/69.073/70.068/71.074). For the fragment information, annotation in MassBank was employed (https://massbank.eu/MassBank/). Peak intensities values were subtracted from the XCMS analysis. Values were normalised by the total sum of features per sample and pareto scaled before statistical analysis. Peak intensities were converted into fold change values using the water treatments for mock and infected plants across the different timepoints. Statistical analysis was performed as described in the *Statistical analyses* section.

### Integration transcriptome and metabolome data

Transcriptomic and metabolomic data were integrated to identify putative correlations. For this, BABA-primed DEGs and BABA-primed putatively identified metabolites at the MS2 level datasets were used for this integration. Analysis was performed using pRocessomics (https://github.com/Valledor/pRocessomics, Escandón et al., 2021) in R v4.3.1 (R_Core_Team, 2023) following the developers’ indications. Data were pre-filtered with a Random Forest for missing values imputation and average intensity for abundance balancing normalisation. After pre-filtering, SPLS was performed to identify the relationships between transcripts (selected as ‘predictor matrix’) and metabolites (selected as ‘response matrix’). Correlation threshold for the network was 0.7. For the visualisation and network export as .png at 360 dpi, Cytoscape 3.10.2 software (https://cytoscape.org/download.html) was used.

### Data availability

All data are available in the different repositories and platforms as follows: R packages and scripts can be found in the public GitHub folder. Transcriptomic data have been deposited in Gene Expression Omnibus. Metabolomic data have been deposited in MetaboLights (Yurekten et al., 2024). Data will be available upon manuscript acceptance.

## Results

### Induced resistance phenotypes after treatment with BABA, JA and SA

To assess chemical-induced resistance, we treated 3-month-old oak seedlings with different elicitors, salicylic acid (SA), jasmonic acid (JA) or β-aminobutyric acid (BABA), followed by infection with the PM causal agent, *Erysiphe alphitoides*, 7 days post elicitor treatment. After inoculation, disease progression was compared against water treated plants. At 14 dpi, JA-treated plants displayed the highest number of colonies (a median of 10 colonies compared to 4 for water), whilst BABA- and SA-treated had fewer (both had a median of zero colonies), however these differences were not statistically significant (Fig. S1A). Resistance phenotypes following elicitor treatment were monitored in both laboratory and field conditions by measuring the Disease Severity Index (DSI). Under laboratory conditions both BABA and SA treatments were associated with significantly lower DSI values at 30 dpi, compared to the water control. Whilst a reduction in DSI was also observed for plants treated with JA, this was not significantly different from the control (Fig. 1A). Under field conditions, a significant reduction in disease severity at 30 dpi was confirmed only for BABA treatment, whereas SA and JA had no significant impact on disease severity when compared to water (Fig. 1B).

**Figure 1.**
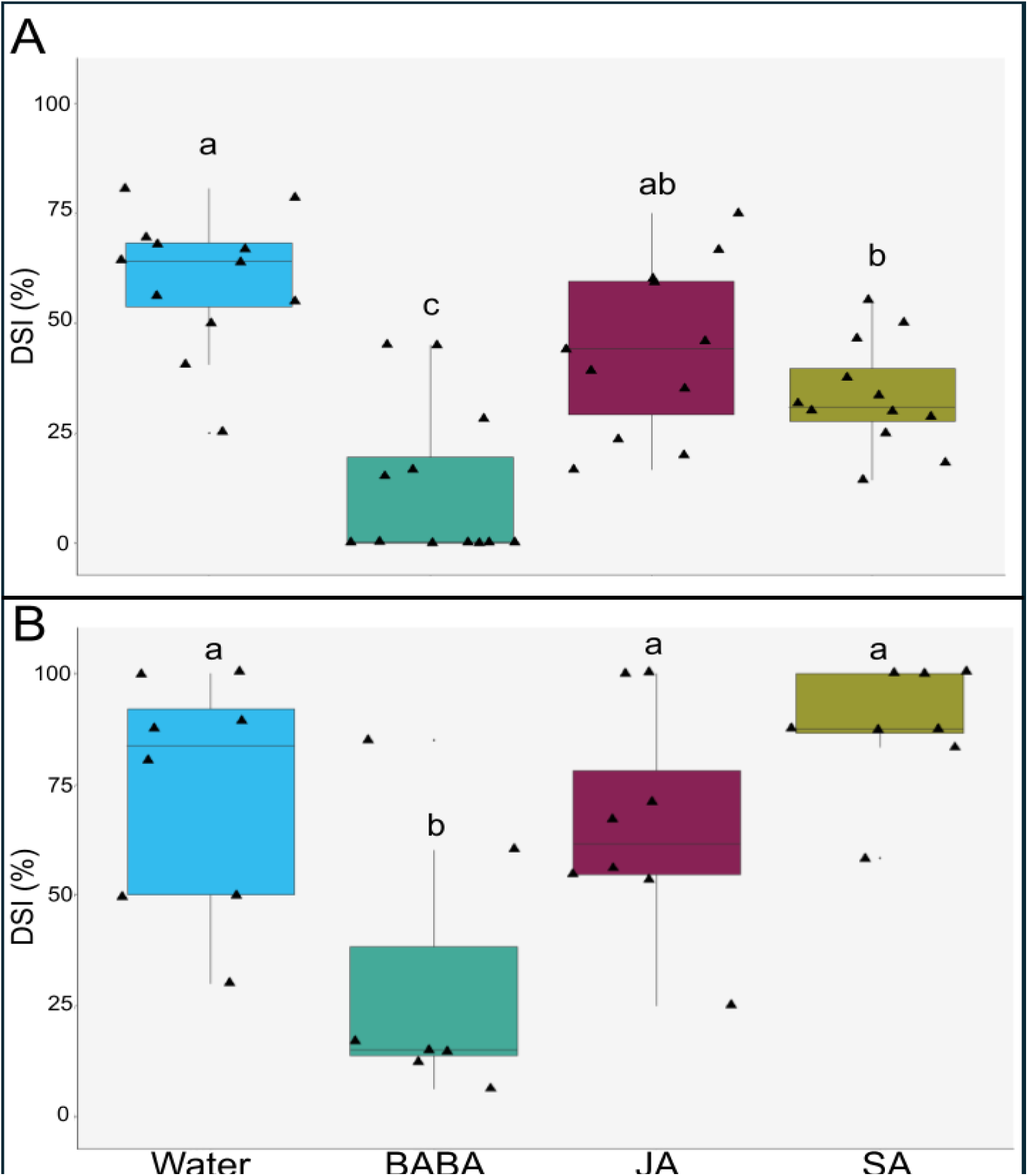
Disease severity after elicitor treatment with BABA, JA and SA. A) Disease Severity Index (DSI, %) under laboratory conditions at 30 dpi. B) DSI (%) under field conditions at 30 dpi. Letters represent statistically significant differences between treatments (one-way ANOVA + Tukey post-hoc test; p ≤ 0.05; n = 11-12 (A) or 7-8 (B)). Black triangles represent biological replicates (seedlings).

### Growth parameters, induced resistance and priming after BABA, JA and SA treatments

All three elicitor treatments had no significant impact on relative growth rate (RGR) for various growth parameters: height (Fig. 2A), stem diameter, leaf length (Fig. S1B-C), or number of leaves (Fig. 2B), when compared to water. Both BABA and JA led to a significant and substantial increase in PR1 expression compared to water in mock-inoculated plants at 1 dpi (left, Fig. 2C). In PM-inoculated plants, BABA and SA led to increased PR1 expression in comparison to water, whilst JA did not. However, these differences were not significant (right, Fig. 2C). Interestingly, SA-treated plants infected with PM showed priming of PR1 expression, as a statistically significant higher expression was observed when compared against water mock, SA mock, or water PM (Fig. 2C). Prior to infection (at 0dpi), all three elicitors had no significant effect on callose deposition compared to water (left, Fig. 2D). Meanwhile, upon PM inoculation (1 dpi) BABA led to a significant increase in the total area of callose compared to all treatments at 0 dpi and 1 dpi (right, Fig. 2D-E). Compared to water, both BABA and SA treated plants displayed a significant increase in the percentage of effective callose (i.e. callose that was able to limit pathogen growth) upon infection, whilst JA had no significant impact (Fig. 2F).

**Figure 2.**
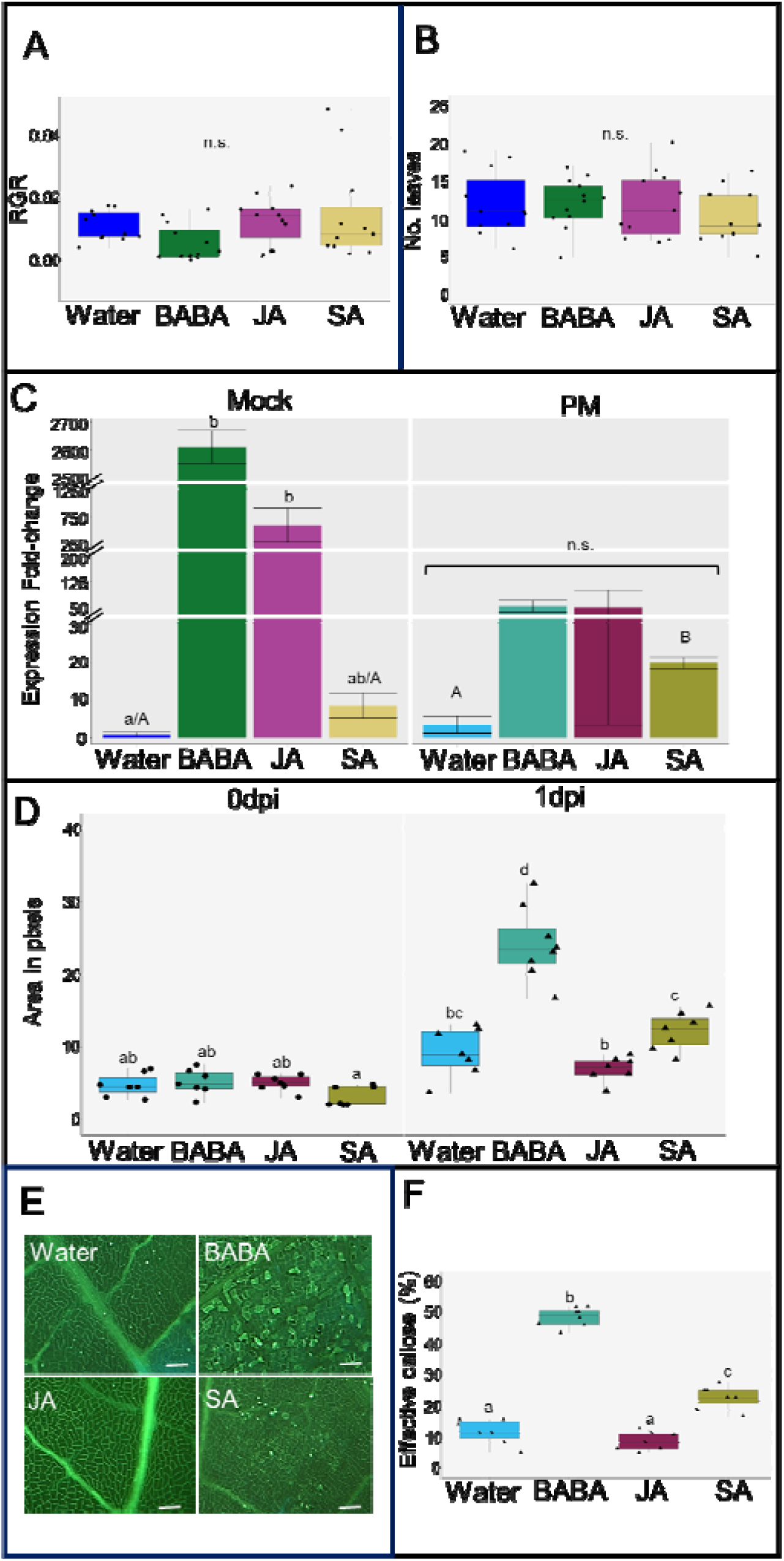
Growth, defence and priming phenotypes after BABA, JA and SA treatments. A) Relative growth rate (RGR) per day of height. B) Number of leaves at t2 (21 days) for mock plants. For both A and B one-way ANOVA was not significant (p>0.05; n = 11-12). C) Fold-change in *PATHOGENESIS RELATED 1* (*PR1*) gene expression for mock and infected plants at 1 dpi. Bar graphs represent the mean and error bars represent the standard error of the mean (SEM). Lowercase letters represent statistically significant differences between treatments within the mock and the PM groups. A Kruskal-Wallis was performed separately for mock (Dunn’s post-hoc test, p ≤ 0.05, n = 2-3) and infected (PM) (not significant, p > 0.05, n = 2-3) plants. Capital letters indicate statistically significant differences between SA and water treatments of both mock and infected (PM) plants (ANOVA + Tukey post-hoc test, p ≤ 0.05, n = 2-3). D) Total callose area in pixels before (0dpi) and after (1dpi) infection. Letters represent statistically significant differences (Welch ANOVA + Dunnett’s T3 post-hoc test, p ≤ 0.05, n = 7-8). E) Representative images of callose deposition at 1 dpi following water, BABA, JA, and SA treatments, respectively. Scale bar = 60μm F) Percentage effective callose in infected plants at 1 dpi. Letters represent statistically significant differences (ANOVA + Tukey post-hoc test; p ≤ 0.05, n = 7-8). Black dots and triangles represent biological replicates (plants) for mock and infected plants, respectively.

### Transcriptome analysis of BABA, JA and SA priming

To further investigate the molecular phenotype induced by elicitor treatments, we analysed the transcriptomes of all conditions by RNA-Seq. Principal component analysis (PCA) was used to assess how the various elicitor treatments differed from each other in mock and infected plants. At 0 dpi, in mock plants, BABA showed clear separation from water and JA, but not SA. Meanwhile, SA only separated from JA. Finally, JA was the only treatment to form a cluster distinct to all other treatments (Fig. 3A). At 1 and 2 dpi, there was some minor separation between mock and PM-infected plants for each elicitor individually (in order of most to least separated: water, SA, BABA, JA) (Fig. 3B-C). For all three timepoints, each component only explained a small proportion of the variance in the data (a maximum of 17% for PC1 at 0 dpi) (Fig. 3A-C). Partial least-squares discriminant analysis (PLS-DA) showed clearer separation of treatments at all timepoints. For 0 dpi, all elicitor treatments formed distinct clusters. Moreover, separation between mock and infected plants for each elicitor individually was slightly clearer at 1 and 2 dpi (Fig. S2).

**Figure 3:**
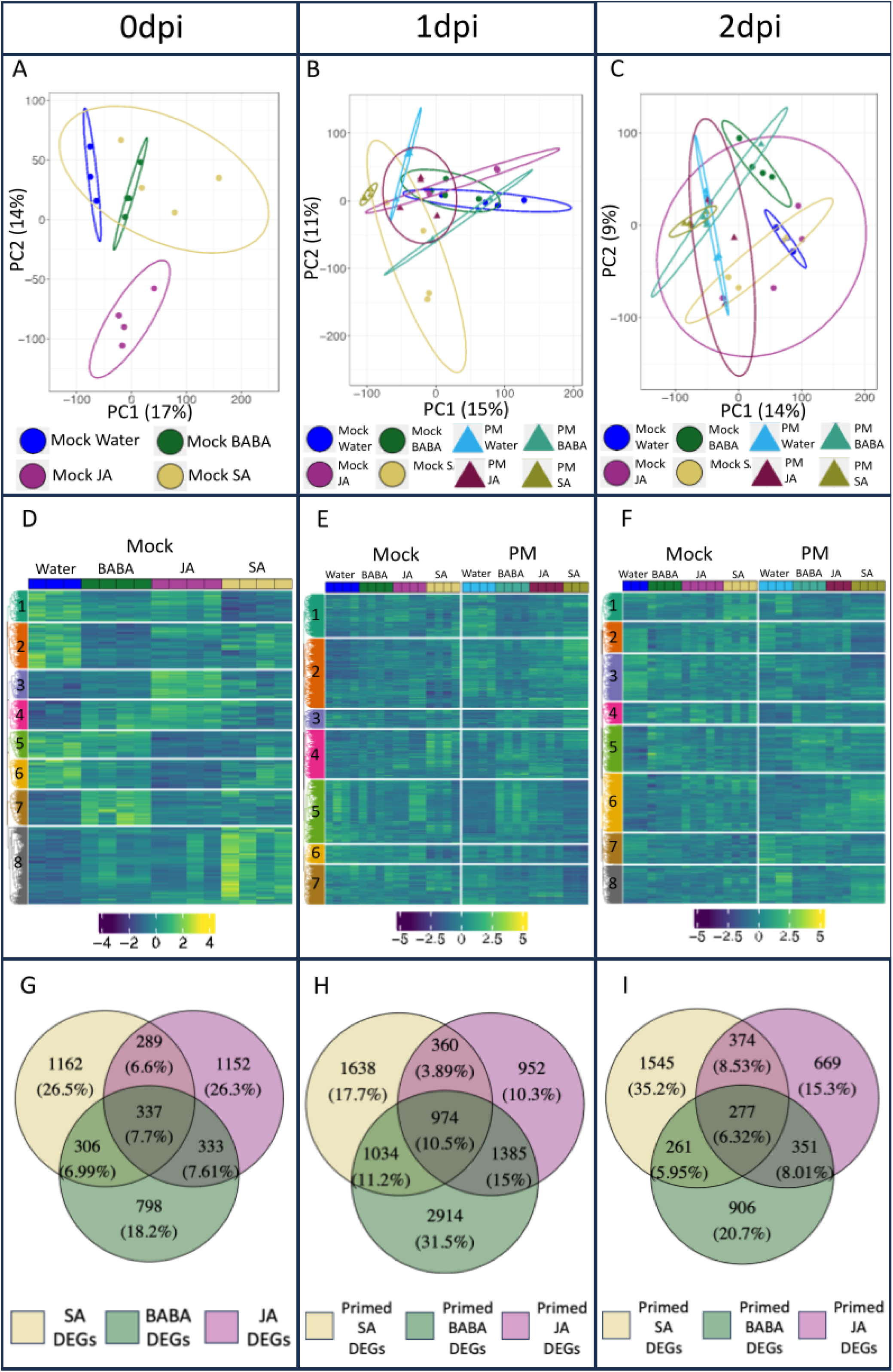
Transcriptome analysis. A) Principal component analysis (PCA) score plot for plants at 0 dpi. B-C) PCA score plots for mock- and PM-infected plants at 1 dpi and 2dpi, respectively. Individual points in PCA plots represent biological replicates (seedlings). D) Heatmap of significant (Wald-test, padj ≤ 0.05) differentially expressed genes (DEGs) for plants at 0 dpi. E-F) Heatmaps of significant (Wald-test, padj ≤ 0.05) DEGs for mock- and PM-infected plants at 1 dpi and 2 dpi. Individual boxes at top of heatmaps represent biological replicates. Numbers represent the identified clusters. G) Venn diagram comparing significant (Wald-test, padj ≤ 0.05) DEGs at 0 dpi. H-I) Venn diagrams comparing significant (Wald-test, padj ≤ 0.05) primed DEGs at 1 dpi and 2 dpi.

Heat-map and hierarchical clustering analysis revealed 8 clusters for 0 and 2 dpi and 7 clusters for 1 dpi (Fig. 3D-F). At 0 dpi (no infection), clusters 7, 8, and 3 corresponded with genes directly upregulated by BABA, SA and JA, respectively. Cluster 4 grouped genes upregulated by BABA and JA whereas clusters 1, 2, 5, and 6 corresponded with genes generally downregulated by specific elicitors. From these, cluster 2 grouped genes that were clearly downregulated by BABA (Fig. 3D). Specific clustering at 1 and 2 dpi is less obvious than at 0 dpi (Fig. 3E-F). At 1 dpi cluster 2 and 7 grouped genes associated with SA and infection, being up- and downregulated, respectively. Cluster 5 corresponded with genes upregulated by BABA upon infection, whilst cluster 1 grouped genes downregulated by BABA upon infection (Fig. 3E). At 2 dpi cluster 6 grouped genes upregulated by SA upon infection, whilst cluster 2 corresponded with genes downregulated by SA upon infection. Cluster 8 corresponded with genes downregulated by BABA and JA (Fig. 3F).

Venn diagrams were used to isolate genes under the control of each elicitor. At 0 dpi, the influence of each elicitor was investigated by comparing their effects with the water treatment. SA and JA had a large effect on the transcriptome with more differentially expressed genes (DEGs) exclusive to these treatments (1162 (26.5%) with SA, 1,152 (26.3%) with JA, compared to 798 (18.2%) with BABA) (Fig. 3G). At 1 dpi and 2 dpi, we isolated the elicitor-specific primed genes: genes that were differentially expressed after elicitor and PM infection only. This isolation was performed by comparing the elicitor + PM treatments to the elicitor + mock treatments in each timepoint. At 1 dpi, genes primed by BABA corresponded with the most extensive transcriptomic changes, with 2,914 (31.5%) DEGs exclusive to this elicitor (Fig. 3H). Meanwhile, at 2 dpi SA triggered the most extensive changes, with 1,545 (35.2%) of DEGs exclusive to this priming elicitor (Fig. 3I). Priming of gene expression by SA was consistent across 1 dpi and 2 dpi as there are only 93 fewer exclusive DEGs at 2 dpi. However, for BABA, much of the transcriptomic changes occurred at 1 dpi and had stopped by 2 dpi as there were 2,008 fewer exclusive DEGs at this later timepoint. Finally, JA had a much more limited impact on differential expression of primed genes with only 952 and 669 genes exclusively differentially expressed at 1 and 2 dpi, respectively (Fig. 3H-I). The highest percentage of overlapping DEGs (15%) occurred at 1 dpi between BABA and JA genes (Fig. 3H). Moreover, the percentage of genes differentially expressed after all elicitor treatments was the highest at this time point (10.5%).

### GO term enrichment analysis

GO term enrichment analysis (Table 1, Tables S1-S6) was performed on the identified primed genes for all the elicitors and timepoints (genes included in Fig. 3H-I). Data input was done using three distinct datasets of primed DEGs: those either up- or downregulated genes (represented as ‘up/down’), those solely upregulated, and those solely downregulated. At 1dpi the Molecular Function (MF) term *heme binding* and Cellular Component (CC) term *membrane* clearly differentiate the three elicitors: in both cases being enriched in downregulated, upregulated, and both up- and downregulated genes for BABA, SA, and JA, respectively. Meanwhile, at 2dpi the MF terms *ADP binding* and *iron ion binding* show a similar pattern of being enriched in different sets of genes for the three elicitors. However, for CC terms at 2dpi and Biological Process (BP) terms at both timepoints there are no GO terms that clearly differentiate all three elicitors in this way. Pair-wise comparison between elicitors unravelled that overall, across both time points there is a larger overlap of GO enriched pathways between BABA and JA (41%), than between BABA and SA (17%) and SA and JA (21%). This overlap between BABA and JA was more pronounced in the BP (52%) and the CC (50%) than in the MF (30%). Therefore, similarly to the overlap of DEGs, BABA and JA appear to perform similarly at a mechanistic level.

**Table 1.** Summary of GO term enrichment analysis of primed DEGs at 1 and 2 dpi. Only those enriched GO terms shared between at least two of the three elicitors are included for clarity. Fisher p-values and the number of genes associated with each term can be found in Tables S1-S6. Up/Down = enriched in data containing all (up- and downregulated) primed DEGs (Wald-test padj ≤ 0.05). Up = enriched in data containing only upregulated primed DEGs (Wald-test padj ≤ 0.05, log2 FC > 1). Down = enriched in data containing only downregulated primed DEGs (Wald-test padj ≤ 0.05, log2 FC < -1). Up + Down = pathways separately enriched in both the datasets containing either upregulated or downregulated primed DEGs. BP = biological process, MF = molecular function, CC = cellular compartment.

| Ontology | GO term | 1dpi |  |  | 2dpi |  |  |
| --- | --- | --- | --- | --- | --- | --- | --- |
|  |  | BABA | JA | SA | BABA | JA | SA |
| BP | plant-type secondary cell wall biogenesis | Up/Down | Up/Down | - | - | - | - |
|  | lignin biosynthetic process | Up/Down | Up/Down | - | - | - | - |
|  | photosynthesis | Up/Down | - | Up/Down | - | - | - |
|  | translation | Up/Down | - | Up/Down | Up/Down | Up/Down | - |
|  | reactive oxygen species metabolic process | - | - | - | Up/Down | Up/Down | - |
|  | rRNA processing | - | - | - | Up/Down | Up/Down | Up/Down |
|  | phosphorelay signal transduction system | - | - | - | Up/Down | Up/Down | Up/Down |
|  | plant-type secondary cell wall biogenesis | Up | Up | - | - | - | - |
|  | lignin biosynthetic process | Up | Up | - | - | - | - |
|  | lignin catabolic process | Up | Up | - | - | - | - |
|  | cellular response to hypoxia | - | Up | Up | - | - | - |
|  | ethylene-activated signaling pathway | - | Up | Up | - | - | - |
|  | regulation of DNA-templated transcription | - | - | - | - | Up | Up |
|  | defense response | Down | Down | - | Up | Up | Up |
|  | defense response to other organism | - | Down | Down | - | - | - |
|  | signal transduction | - | Down | Down | - | - | - |
|  | response to molecule of bacterial origin | - | - | - | Down | Down | - |
|  | protein phosphorylation | - | - | - | Down | Down | - |
|  | positive regulation of defense response | - | - | - | Down | Down | - |
|  | plant-type hypersensitive response | - | - | - | Down | Down | - |
|  | reproductive process | - | - | - | Down | Down | - |
|  | cellular nitrogen compound metabolic process | - | - | - | Down | - | Down |
|  | recognition of pollen | - | - | - | - | Down | Down |
| MF | fatty acid synthase activity | Up/Down | Up/Down | - | - | - | - |
|  | transcription cis-regulatory region binding | Up/Down | Up/Down | - | - | - | - |
|  | water channel activity | Up/Down | Up/Down | - | - | - | - |
|  | mRNA binding | Up/Down | - | Up/Down | - | Up/Down | Up/Down |
|  | structural constituent of ribosome | Up/Down | - | Up/Down | Up/Down | Up/Down | - |
|  | glucosyltransferase activity | - | Up/Down | Up/Down | - | - | - |
|  | monooxygenase activity | - | - | - | Up/Down | Up/Down | - |
|  | oxidoreductase activity, acting on paired donors, with incorporation or reduction of molecular oxygen | - | - | - | Up/Down | Up/Down | - |
|  | DNA-binding transcription factor activity | - | - | - | Up/Down | - | Up/Down |
|  | translation initiation factor activity | - | - | - | - | Up/Down | Up/Down |
|  | iron ion binding | - | - | - | Up/Down | Up/Down | Up/Down |
|  | microtubule binding | Up | Up | - | - | - | - |
|  | hydroquinone:oxygen oxidoreductase activity | Up | Up | - | - | - | - |
|  | transcription cis-regulatory region binding | Up | Up | - | - | - | - |
|  | polysaccharide binding | - | Up | Up | - | - | - |
|  | oxidoreductase activity, acting on CH-OH group of donors | - | Up | Up | - | - | - |
|  | heme binding | Down | Up+Down | Up | Up+Down | - | Up+Down |
|  | ADP binding | Down | Down | Down | Up+Down | Down | Up |
|  | NADP+ nucleosidase activity | - | Down | Down | Up | - | Up |
|  | NAD+ nucleosidase activity | Down | Down | Down | Up | - | Up |
|  | NAD+ nucleotidase, cyclic ADP-ribose generating | - | Down | Down | Up | - | Up |
|  | iron ion binding | Down | Down | Down | Up+Down | Down | Up+Down |
|  | monooxygenase activity | Down | Down | Down | - | Down | Down |
|  | oxidoreductase activity, acting on paired donors, with incorporation or reduction of molecular oxygen | Down | Down | Down | - | Down | Down |
|  | ATP binding | Down | Down | Down | Down | Down | - |
|  | protein serine kinase activity | - | - | - | Down | Down | - |
|  | protein binding | - | - | - | Down | Down | - |
|  | ubiquitin protein ligase binding | - | - | - | Down | - | Down |
|  | carbohydrate binding | - | - | - | - | Down | Down |
| CC | chloroplast membrane | Up/Down | Up/Down | - | - | Up/Down | Up/Down |
|  | chloroplast thylakoid membrane | Up/Down | - | Up/Down | - | - | - |
|  | cytosol | Up/Down | - | Up/Down | Up/Down | Up/Down | Up/Down |
|  | chloroplast stroma | Up/Down | Up/Down | Up/Down | - | - | - |
|  | organelle subcompartment | Up/Down | Up/Down | Up/Down | - | - | - |
|  | chloroplast envelope | Up/Down | Up/Down | Up/Down | - | - | - |
|  | cytosolic large ribosomal subunit | - | - | - | Up/Down | Up/Down | - |
|  | ribonucleoprotein complex | - | - | - | Up/Down | Up/Down | - |
|  | extracellular region | Up | Up | - | - | - | - |
|  | microtubule | Up | Up | - | - | - | - |
|  | apoplast | Up | Up | - | - | - | - |
|  | side of membrane | Up | Up | - | - | - | - |
|  | plant-type cell wall | Up | Up | - | Up | - | Up |
|  | cell wall | Up | Up | - | - | - | - |
|  | membrane | Down | Up+Down | Up | - | - | - |
|  | plasma membrane | Down | Down | - | Down | Down | Down |
|  | vacuolar membrane | Down | Down | - | - | - | - |
|  | chloroplast membrane | Down | - | Down | - | - | - |

### Metabolome analysis of BABA, JA and SA priming

Metabolome analysis was performed to further investigate the mechanisms of induced resistance and priming of the three elicitors. PCA showed little to no separation between various elicitor treatments in mock and infected plants across all timepoints (Fig. 4 A-C). Again, PLS-DA showed clearer separation of treatments at all timepoints. At 0 dpi, each treatment showed somewhat distinct clusters, although there remained at least some overlap. Moreover, there was at least some separation between treatments at 1 and 2 dpi, with BABA and water being most different from each other (Fig. S3).

**Figure 4:**
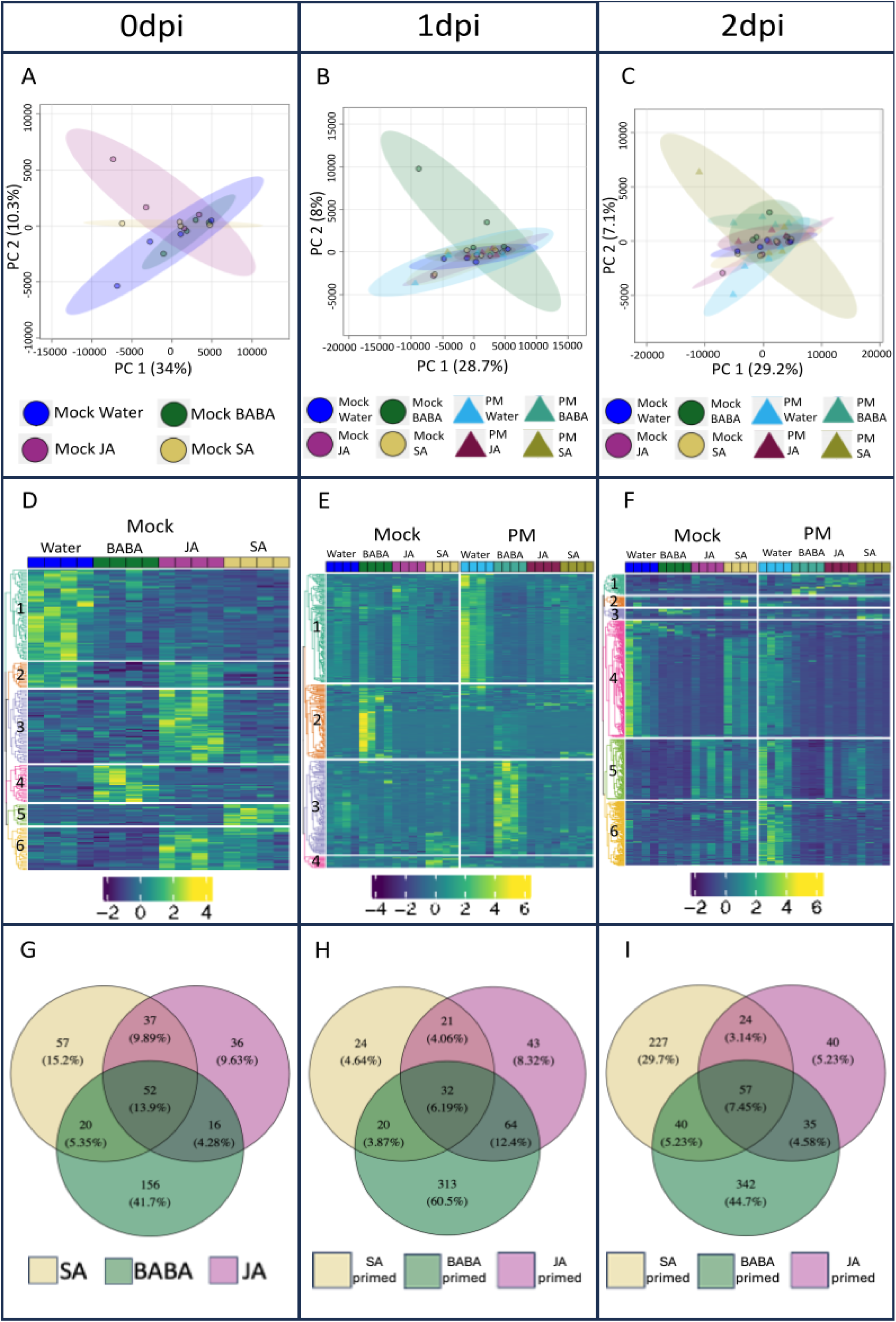
Metabolome analysis. A) Principal component analysis (PCA) scores plot for mock plants at 0 dpi. B-C) PCA scores plots for mock and infected plants at 1 dpi and 2dpi. Individual points in PCA plots represent biological replicates. D) Heatmap of significant (ANOVA, p ≤ 0.01) metabolites for mock plants at 0 dpi. E-F) Heatmaps of significant (ANOVA, p ≤ 0.01) metabolites for mock and infected plants at 1 dpi and 2 dpi. Individual boxes at top of heatmaps represent biological replicates. G) Venn diagram comparing significant (ANOVA, p ≤ 0.05, log2 FC ≥2 or ≤-2) metabolites at 0 dpi. H-I) Venn diagrams comparing significant (ANOVA, p ≤ 0.05, log2 FC ≥2 or ≤-2) primed metabolites at 1 dpi and 2 dpi.

Heat-map and hierarchical clustering analysis revealed 6 clusters for 0 and 2 dpi and 4 clusters for 1 dpi (Fig. 4D-F). At 0 dpi (no infection), cluster 1 was associated with metabolites downregulated by all elicitors. Cluster 2 grouped metabolites downregulated by BABA and SA. Meanwhile, clusters 3, 4, and 5 corresponded with metabolites directly upregulated by JA, BABA and SA, respectively. Finally, cluster 6 was upregulated by JA and SA (Fig. 4D). At 1 dpi, cluster 1 grouped metabolites downregulated by all elicitors upon infection. Clusters 2 and 3 were primarily associated with metabolites upregulated by BABA in mock or infected (PM) plants, respectively. Cluster 4 corresponded to metabolites upregulated by SA in mock plants (Fig. 4E). At 2 dpi, cluster 1 grouped metabolites upregulated by BABA and JA upon infection. Meanwhile, clusters 2 and 3 corresponded with metabolites upregulated by SA in mock and infected plants, respectively. Cluster 5 was primarily associated with metabolites downregulated by BABA, particularly upon infection. Finally, clusters 4 and 6 grouped metabolites downregulated by all three elicitors upon infection (Fig. 4F).

Venn diagrams were used to isolate metabolites under the control of each elicitor. At 0 dpi, BABA had the largest effect on the metabolome with 156 (41.7%) significant metabolites exclusive to this treatment (Fig. 4G). BABA also had the largest effect on primed metabolites at 1 and 2 dpi: 313 (60.5%) and 342 (44.7%) metabolites are exclusive to BABA-induced priming, respectively. Meanwhile, SA only triggered large-scale metabolomic changes at 2 dpi, with 227 (29.7%) metabolites being exclusive to SA-induced priming at this time-point (compared to only 24 (4.64%) at 1 dpi) (Fig. 4H-I). JA had a limited impact on the metabolome at all timepoints, with only 36, 43, and 40 metabolites at 0, 1, and 2 dpi, respectively. The highest percentage of overlapping metabolites between two elicitors occurred at 1 dpi between BABA and JA (12.4%). Moreover, the percentage of metabolites differentially accumulated after all elicitor treatments was the highest at 0 dpi (13.9%) (Fig. 4G-I). Therefore, BABA was the elicitor that triggered the largest changes at a metabolomic level.

The metabolome data was used to investigate patterns of defence hormone accumulation after elicitor treatments. From the four hormones tested (i.e. SA, JA, JA-Val and JA-Ile), only SA and JA-Val were found in the samples. For JA-Val, at 0 dpi only JA-treated plants displayed a significant increase compared to the other treatments (Fig. S4A), whilst at 1 dpi (Fig. S4C) and 2 dpi (Fig. S4E) no significant differences in the levels of JA-Val were observed between the different treatments. For SA at 0 dpi, both SA and JA treated plants had higher levels of SA compared to water and BABA treatments, however this difference was not significant (Fig. S4B). A significant increase in the levels of SA was observed at 1 dpi for SA treated plants compared to the other treatments (Fig. S4D); this pattern had disappeared at 2dpi with no significant differences in the levels of SA between elicitor treatments (Fig. S4F). BABA treatments were not associated with changes in SA or JA-Val levels at any time point (Fig. S4).

Primed masses were selected from the Venn Diagrams (Fig. 4H-I) as specific for BABA, SA and JA at 1 and 2 dpi (i.e. masses present at any or both timepoints). A total of 569 BABA-, 205 SA- and 43 JA-specific masses were putatively annotated by using KEGG *Populus trichocarpa* database and libraries from Pastor’s group (Manresa-Grao et al., 2022; Manresa-Grao et al., 2024). A total of 88 metabolites at level of confidence 3 (MS1 and MS2) were identified (Table S7). From these 88 identified metabolites, 55 were specific to the BABA treatment, 28 to the SA treatment and 5 to the JA (Table S7). In addition, pathway enrichment analysis was performed with Marvis using the identified metabolites. Considering the low numbers, no enrichment was found for SA-primed or JA-primed metabolites. For the identified BABA-primed metabolites, statistically significant enrichment was found for Alkaloids (14 metabolites), Indole derivatives (three metabolites), Lignans (five metabolites), and Phenylpropanoids (six metabolites) (Table 2). At 1 dpi, all except four of these significant metabolites were primed, with most of these being upregulated (15) rather than downregulated (8). Half of the phenylpropanoids were upregulated (three up, two down, one not primed). Most of the alkaloids (nine up, two down, three not primed) and indoles (two up, one down) were upregulated, whilst most of the lignans were downregulated (one up, four down). At 2 dpi whilst the same number of significant metabolites were primed, unlike at 1 dpi, most of them were downregulated (20) rather than upregulated (four). All lignans and indoles were downregulated, as were most of the alkaloids (three up, eight down, three not primed) and phenylpropanoids (four down, one up, one not primed).

**Table 2.**
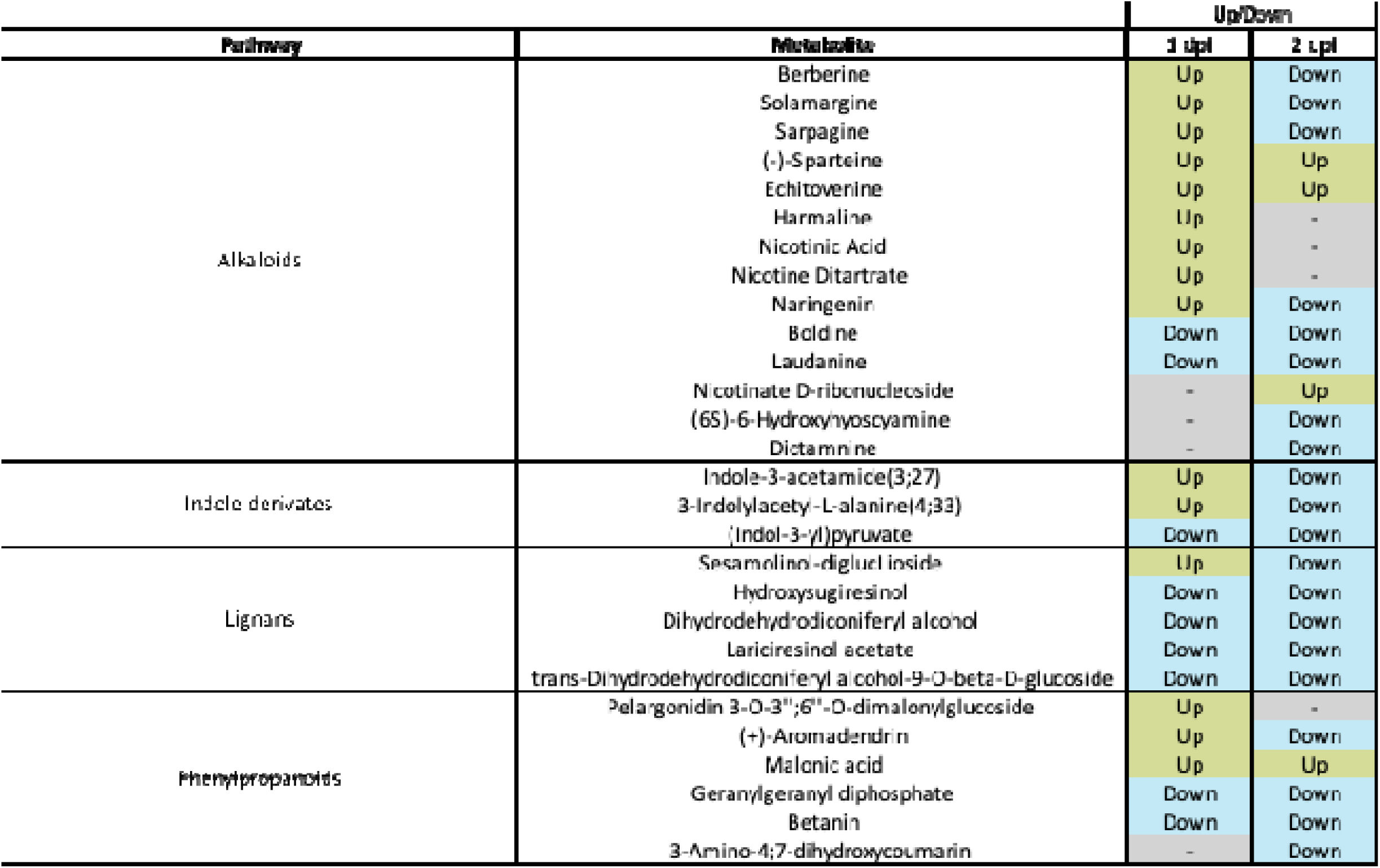
Summary of pathways enriched for BABA priming and the identified metabolites associated to those pathways. Up = metabolites with a fold-change > 2 compared to mock water at that time point. Down = metabolites with a fold-change < 0.5 compared to mock water at that time point.

### Transcriptome and metabolome integration

To further mine the omics results, transcriptomic and metabolomic datasets were integrated. From the transcriptome data, genes associated with priming by BABA were selected: 2914 genes corresponding to 1 dpi and 906 genes corresponding to 2 dpi (Fig. 3H-I), which produced a list of 3648 unique genes across the two timepoints (Table S8). From the metabolome data, the 55 metabolites putatively identified for BABA priming were used in this analysis (Table S7).

This analysis unravelled that only six genes were correlated to only one metabolite: benzene (Fig. 5). Genes were annotated using blast at the level of the *Quercus* genus, which included annotations from *Q. robur*, *Q. suber* and *Q. lobata* (Table S9). *UBIQUITIN FUSION DEGRADATION PROTEIN 1* and *HYDROXYACYL GLUTATHIONE HYDROLASE 2* had the strongest negative correlation (<-0.7) followed by *E3 UBIQUITIN-PROTEIN LIGASE and BETA-CATENIN-LIKE PROTEIN 1* with a negative correlation (<-0.5). *TBC1 DOMAIN FAMILY MEMBER 15* and *UPSTREAM ELEMENT-BINDING PROTEIN 1-LIKE* showed a positive correlation (>0.4).

**Figure 5:**
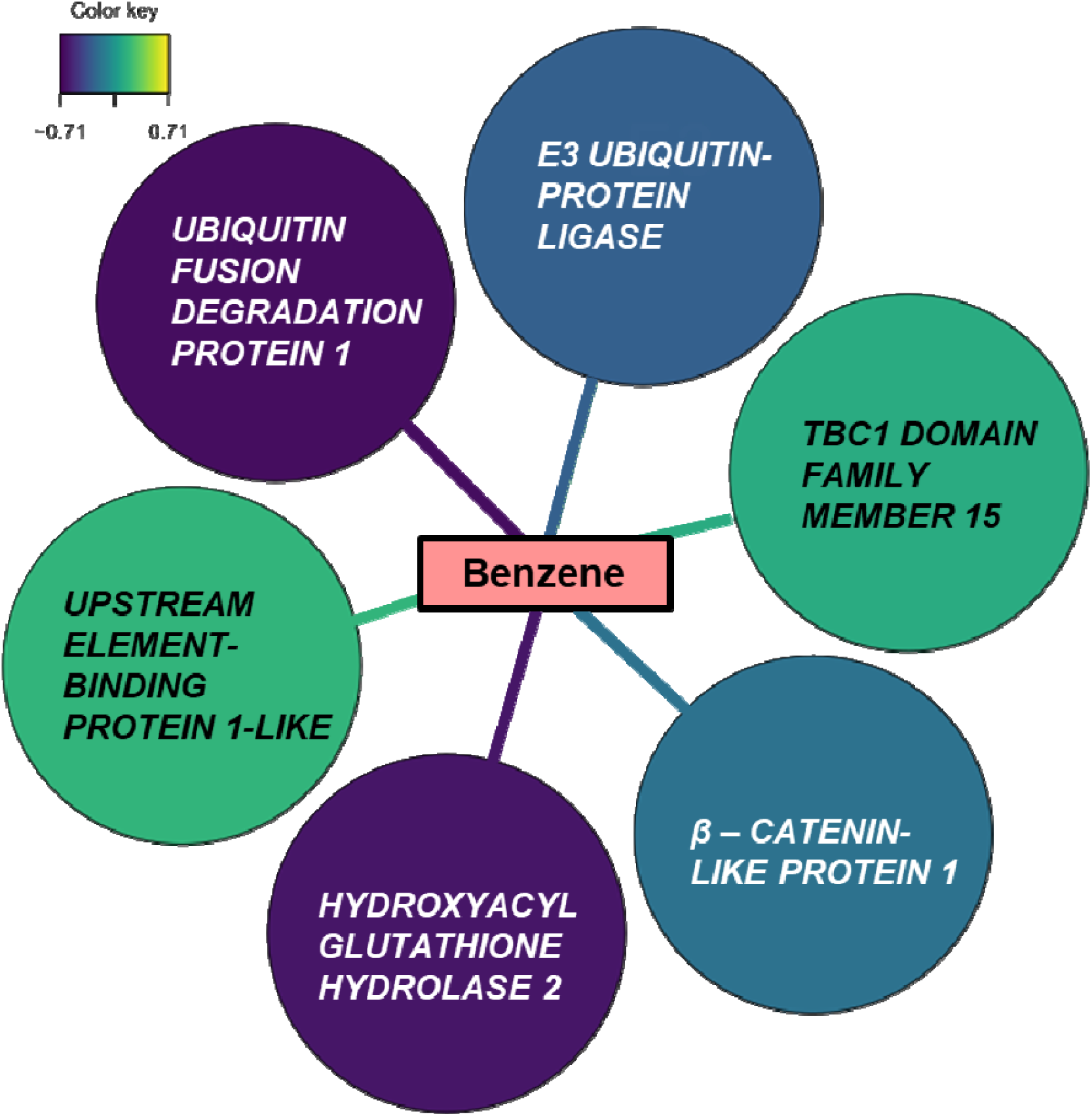
Integration analysis. Transcriptome and metabolome dataset were integrated using a correlation threshold of 0.7 with a confidence of 0.9. Correlation is indicated with the colour of the arrows connecting Transcripts and Metabolite, where purple indicates negative correlation and yellow indicates positive.

## Discussion

Here, we have provided valuable insight into defence priming in oak that may contribute to sustainable strategies to control PM, a major bottleneck for oak woodland regeneration. We have investigated the specific mechanisms of IR and priming of three well-characterised elicitors in the oak seedling-PM pathosystem. We have demonstrated that BABA and SA lead to enhanced resistance to PM in oak seedlings. Whilst SA only enhanced resistance in laboratory conditions, BABA had a consistent impact in both laboratory and field conditions (Fig. 1). It is perhaps to be expected that SA is less effective at increasing resistance, because BABA is known to be especially effective at inducing priming by acting on multiple pathways (Cohen et al., 2016). Meanwhile, JA had no significant impact on resistance in both conditions (Fig. 1). This likely reflects the fact that powdery mildews are biotrophic pathogens; and SA, but not JA, signalling is commonly involved in defending against these species (Glazebrook, 2005; Shigenaga et al., 2017). However, it was notable that JA did not enhance susceptibility either, indicating that the reported antagonism between SA and JA in herbaceous species (Li et al., 2019a) may not be expressed in this pathosystem. This was also studied by quantification of defence hormones: accumulation of JA-Val and SA remains similar in both JA- and SA-treated plants at all timepoints, which further suggests that SA/JA defence crosstalk is not expressed in oak seedlings. This is supported by a lack of antagonism seen in other woody plants such as *Populus* (Ullah et al., 2022) and in *Citrus aurantium* (Agut et al., 2014). This result is somewhat expected due to the many different exposures that trees can experience at one specific time in their lives. Specifically, oak seedlings would not benefit from a prioritisation of defence mechanisms as it would result in the compromise of specific pathways that could damage their growth and development. Overall, of the three elicitor treatments, only BABA consistently enhances resistance to PM in a manner applicable to reducing its impact on woodland regeneration.

We have shown that certain chemical elicitors (i.e., BABA and SA) increase resistance to PM in oak, however it is necessary to determine if this is due to direct induction or priming of plant defences. For example, increased resistance to *Ips typographus* beetles in Norway spruce (*Picea abies*) can occur due to either prolonged direct induction of defences following fungal infection or priming of defence following MeJA treatment (Mageroy et al., 2020a). Moreover, Bengtsson et al., 2014, showed that in potato BABA-IR is associated with direct induction of defences rather than priming. Direct induction of resistance involves a trade-off between the activation of defence responses and a reduction in plant growth (He et al., 2022). For instance, studies in various plants (e.g., Arabidopsis, rice, tobacco, and tomato) have demonstrated that BABA, JA, and SA treatments, particularly those concentrations required for direct induction of defence, can all reduce growth (Luna et al., 2016; Bhavanam and Stout, 2021; Li et al., 2022). However, as demonstrated for both relative growth rates and number of leaves, none of the treatments resulted in a growth reduction (Fig. 2A-B). This confirms previous findings in oak where no impacts on growth were observed after BABA treatment (Sanchez-Lucas et al., 2023). Moreover, both SA and JA treatment had no significant impact on growth, suggesting that these hormones are unlikely to be directly inducing defence mechanisms. Interestingly when looking at the targeted hormones identified in the oak seedlings (Fig. S4B, D and F), unlike in other plant species BABA did trigger direct accumulation of SA (Li et al., 2019b; Kim and Kang, 2023) which can compromise growth. Since BABA does not lead to SA accumulation its mode of action in oak may be independent from SA signalling.

A phenotype commonly associated with SA-mediated defences and the SAR priming response is increased *PATHOGENESIS-RELATED PROTEIN 1 (PR1)* expression (Backer et al., 2019). Whilst no significant differences were observed when comparing all treatments, comparison of just water and SA for both mock and infected plants demonstrated that SA does indeed prime *PR1* expression in oak seedlings (Fig. 2C). Interestingly, despite no consequences for growth, it was clear that both BABA and JA directly induce, rather than prime, *PR1* expression. Moreover, the increase in resistance associated with BABA-IR has been linked to ABA-dependent callose deposition which acts to restrict pathogen invasion and spread (Schwarzenbacher et al., 2020). SA and the associated priming response of SAR has also been linked to callose deposition (Wang et al., 2021). Here, we demonstrate this phenomenon occurs in this pathosystem, at least upon initial infection (1 dpi). Whilst only BABA led to a significant increase in the total area of callose; both BABA and SA triggered an increase in the percentage of effective callose (callose that has managed to limit pathogen growth). It was notable that in terms of effective callose, BABA led to roughly twice the amount as SA, which may help explain the more effective and consistent resistance observed for BABA. Therefore, these data demonstrate that whilst SA primes different defence signalling pathways, the effects of BABA may be the result of a combination of direct defence gene induction and priming of callose.

Overall, since we have demonstrated that no growth reduction occurs and that SA and BABA prime *PR1* expression and callose deposition, respectively, observed differences in induced resistance between treatments, may be due to expression of priming of defence. This is because priming, by triggering a transient expression of defence mechanisms (Cooper and Ton, 2022), is thought to be less costly (van Hulten et al., 2006) than direct induction of defences. To assess how this priming response is connected to altered gene expression, an untargeted transcriptome analysis was performed. It was clear that BABA-priming of the transcriptome is more rapid and short-term (the response peaks at 1 dpi and has reduced by 2 dpi) than SA-priming (there is a high number of DEGs at 1 and 2 dpi). Venn diagrams unravelled a greater overlap between BABA primed genes and those primed by SA and JA at 1 dpi than at 2 dpi. GO enrichment analysis (Table 1, Supplementary Tables 1-6) revealed that all three elicitors, but particularly BABA, enrich more GO terms at 1 dpi (45) compared to 2 dpi (36) which may reflect the transient expression of defence mechanisms commonly associated with priming (Cooper and Ton, 2022). In the case of BABA and SA, variations in oak seedling resistance to PM following elicitor treatment can be at least partially explained by differences in specific transcriptomic changes triggered following priming by the different elicitors. Interestingly, at both timepoints, Venn diagrams showed that the overlap between BABA with JA was greater than with SA. Whilst BABA-IR has often been linked to SA signalling (Siegrist et al., 2000; Zimmerli et al., 2000; Ton et al., 2005; Eschen-Lippold et al., 2010), particularly in the case of the GO enrichment analysis, overlap between these two elicitors appears limited other than enriching pathways involved in photosynthesis and chloroplast metabolism (Table 1). Instead, at this level we observe again a larger overlap between BABA and JA, where pathways involved in lignans, and defence responses are enriched (Table 1). This is interesting because even though it has been shown previously that JA plays a role in BABA-IR in grapevine (Hamiduzzaman et al., 2005) and tomato (Zapletalová et al., 2023), in our system we do not observe JA-induced resistance against PM. It remains to be seen whether this link between BABA-IR and JA is consistent across other stresses or whether, similar to findings in Arabidopsis (Zimmerli et al., 2000; Ton and Mauch-Mani, 2004; Jakab et al., 2005), the exact mechanism behind BABA-IR varies depending on the specific stress encountered.

To properly understand a complex response like priming it is necessary to combine multiple omics approaches. For example, ISR in Arabidopsis has been linked to large-scale metabolome and limited transcriptome changes in some cases (Brotman et al., 2012) but large-scale transcriptome and limited metabolome changes in others (van de Mortel et al., 2012). Therefore, this study also looked at whether elicitor induced resistance was due to priming of metabolites using an untargeted metabolome analysis. Unlike with the transcriptome, differences between elicitor treatments and mock and infected plants were unclear following PCA at all timepoints (Fig. 4A-C). However, PLS-DA did reveal at least some minor differences between elicitors at 0 dpi and when comparing mock and infected plants (Fig. S5). More limited separation may indicate that changes in the metabolome contribute less to priming in oak than transcriptome changes. However, heatmaps and clustering analysis did indicate that metabolomic changes do play at least some role. For example, at 0 dpi each elicitor was associated with at least one specific upregulated cluster (Fig. 4D). Moreover, at 1 and 2 dpi, especially for BABA, it was clear that each elicitor was associated with specific clusters of metabolites (Fig. 4E-F). This suggests that differences in resistance following elicitor treatment prior to infection are linked to at least some changes in metabolite expression. Venn diagrams revealed that BABA has the largest impact on the metabolite profile of the seedlings at all timepoints (Fig. 4G-I). Similarly to observations in the transcriptome, BABA primes metabolome changes earlier than SA, with the response in BABA being clearly visible at 1 dpi but only becoming visible for SA at 2 dpi. The larger-scale impact of BABA on the metabolome may be one of the reasons it is more consistent and effective at increasing resistance to PM.

To further explore the mechanisms of priming by all the elicitors, we identified metabolites at a level of confidence 3 (MS1 and MS2). A higher number of metabolites were identified for BABA than for SA and JA, however, on a global level, the identification percentage was similar (10-15%). Most of the metabolites identified (Table S7) have been linked to plant defence mechanisms, primarily in redox metabolism for the metabolites associated with BABA priming. For instance, Nicotinic acid has been demonstrated to enhance resistance in spruce seedlings against insects (Berglund et al., 2015). Moreover, Pelargonidin 3-O-3’’;6’’-O-dimalonylglucoside and Betanin are both phenylpropanoids with pigment properties in plant tissues (Esatbeyoglu et al., 2015; Wang et al., 2023) with a role in plant defence. Naringenin has been also described as a plant defence driver, for example conferring resistance to *Phytophthora* in tobacco (Sun et al., 2022) via SA-pathway induction, increasing the expression of *PR*1. Interestingly, JA was associated with altered accumulation of the glucose conjugate of ABA (i.e. ABA glucose ester), which has been associated with priming of callose (Schwarzenbacher et al., 2020). Some of the identified compounds such as berberine and solamargine have also been shown to display effective toxicity against herbivores and pathogens (Thawabteh et al., 2019). To identify pathways, enrichment analysis was performed in these compounds. Due to low numbers, pathway enrichment analysis failed to identify any pathways associated with SA and JA. In contrast, several pathways were identified for the BABA-priming metabolites: Alkaloids, Indole derivatives, Lignans and Phenylpropanoids (Table 2). The role of these pathways does not come as a surprise; BABA-induced resistance has been linked to activation of these defence pathways in other species (Gamir et al., 2012; Wilkinson et al., 2018; Yao et al., 2020; Zapletalová et al., 2023). Future research will focus on enhancing our understanding on the role of the identified metabolites and pathways associated with BABA and will further explore the responses primed by SA and JA.

The integration of transcriptome and metabolome dataset showed only one node around the benzene (Fig. 5). Although benzene is not naturally present in plants on its own, benzene is the basic structure of multiple secondary metabolites with an aromatic ring as terpenoids, alkaloids and phenylpropanoids (Greger, 2017; Al-Khayri et al., 2023; Siddiqui et al., 2024). Its abundance can be altered by the synthesis/degradation of these plant defence compounds. Interestingly, all these pathways associated with benzene are the ones enriched in BABA-treated plants. Our analysis identified six genes linked to Benzene (Table S9), which have been associated with plant immunity in other plant species. For instance, *HYDROXYACYL GLUTATHIONE HYDROLASE 2* (also known as *GLYOXALASE II*) functions to detoxify/breakdown methylglyoxal, overaccumulation of which is particularly common during both biotic and abiotic stress responses in plants (Li, 2016). We also identify a *E3 UBIQUITIN-PROTEIN LIGASE (AR17)* with a RING domain. E3 ligases perform ubiquitination to regulate many aspects of plant immunity, including pathogen recognition, hormone signalling (including SA and JA pathways), and activation of defence responses (Marino et al., 2012; Kelley, 2018).

They have also been linked to BABA-IR in peach fruit (Li et al., 2020). *ß-CATENIN-LIKE PROTEINS* are homologous to ARM-type proteins in plants which have been linked to light/gibberellin signalling, self-incompatibility, trichome development, ABA signalling and receptor-kinase signalling (Coates et al., 2006). Therefore, the identified genes are part of complexes with common functions in mediating the ubiquitination and subsequent degradation of target proteins, which can influence various signalling pathways. Further research will focus on understanding the mode of action of these proteins in BABA-induced priming in oak.

This study has characterised chemical-induced priming during infection of oak seedlings with PM using both transcriptomics and metabolomics analysis. This not only provides valuable insight into the priming response in forest trees but may directly contribute to the development of sustainable strategies that reduce the impact of PM and improve the effectiveness of oak woodland regeneration.

## Acknowledgements

We are extremely grateful to the Birmingham Institute of Forest Research (BIFoR) which provided the space for field work experiments described in this publication. We thank the SCIC of the Universitat Jaume I for technical assistance in metabolomics. The work was funded by the JABBS foundation grant ‘Resistance strategies of oak trees in the arms race with pathogens’ to E.L and RSL and the UKRI grant “MEMBRA” NE/V021346/1 to EL and MC. The work of JLB is supported by the University of Birmingham Midlands Integrative Biosciences Training Partnership (MIBTP) grant BB/T00746X/1. The work of VP is supported by the Valencian grant CIACO/2021/092.

---

**Figure S1:**
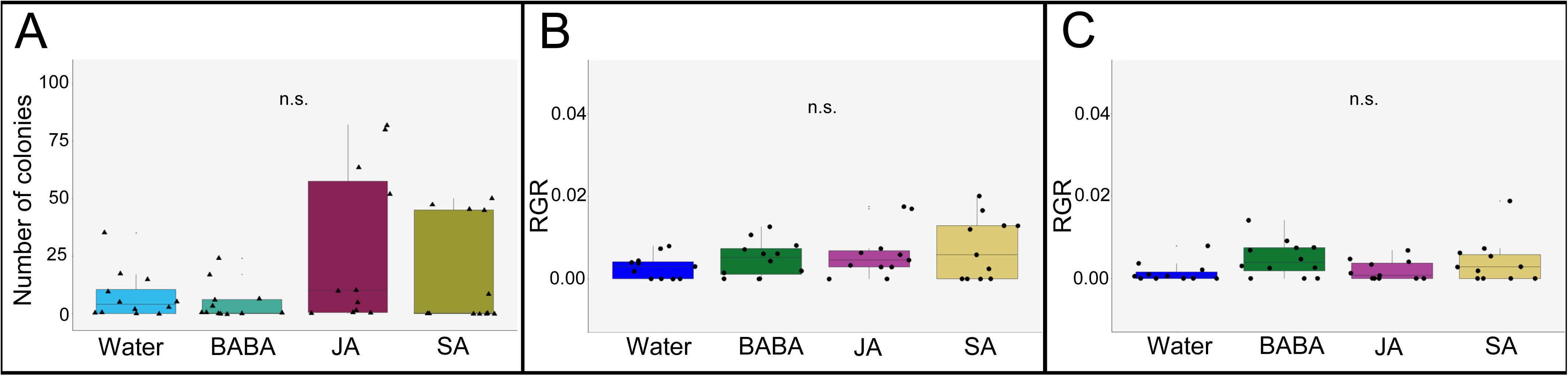
Growth, disease and resistance phenotypes. A) Number of colonies at 14 dpi. B) Relative growth rate (RGR) per day for diameter. C) Relative growth rate (RGR) per day for leaf length. For A,B and C Kruskal-Wallis was not significant (p > 0.05, n = 11-12).

**Figure S2:**
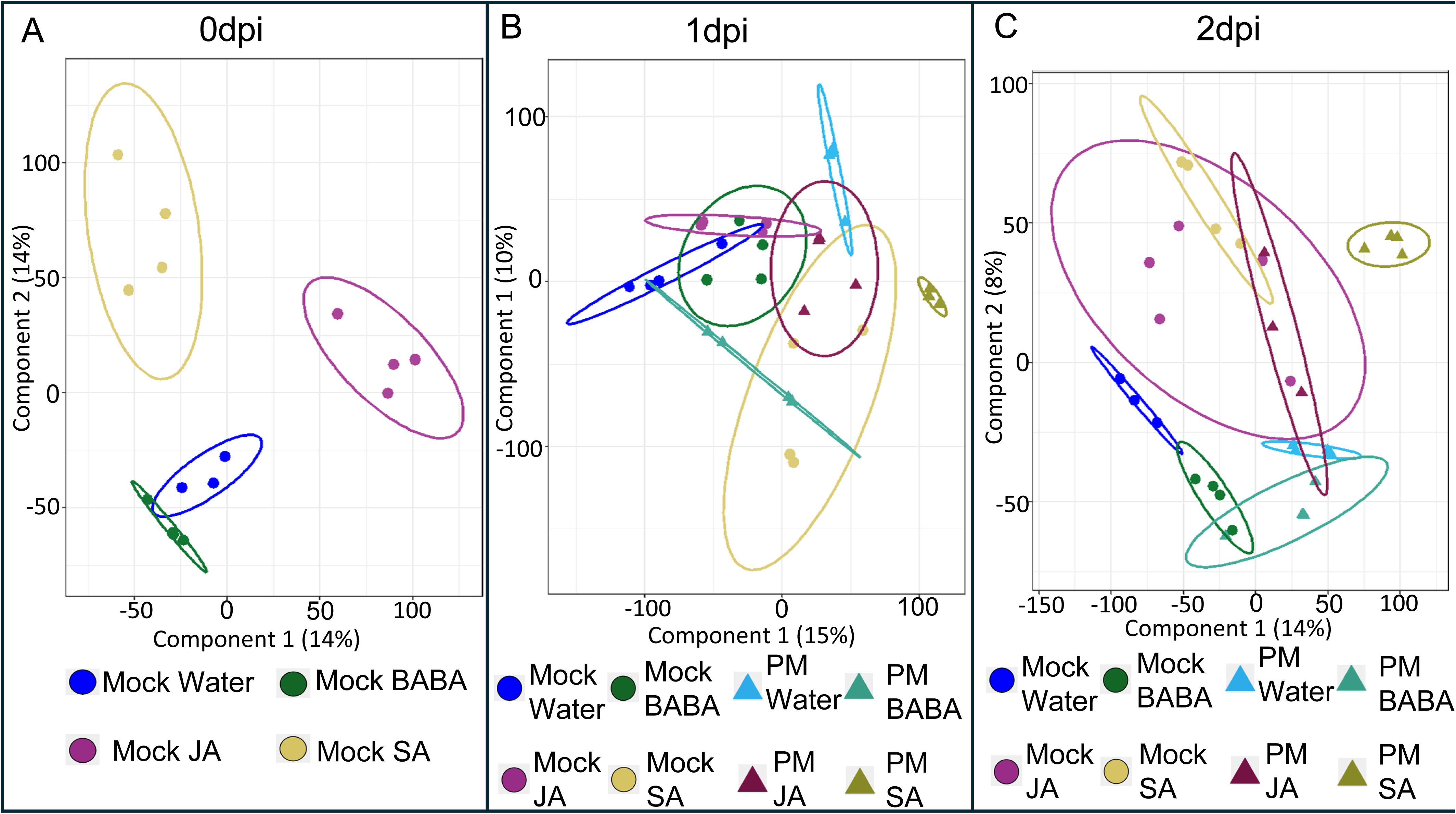
Transcriptome PLS-DA plots. A) Partial Least-Squares Discriminant Analysis (PLS-DA) score plot for mock plants at 0 dpi. B-C) PLS-DA score plots for mock and infected plants at 1 dpi and 2dpi, respectively. Individual points in PLS-DA plots represent biological replicates (plants).

**Figure S3:**
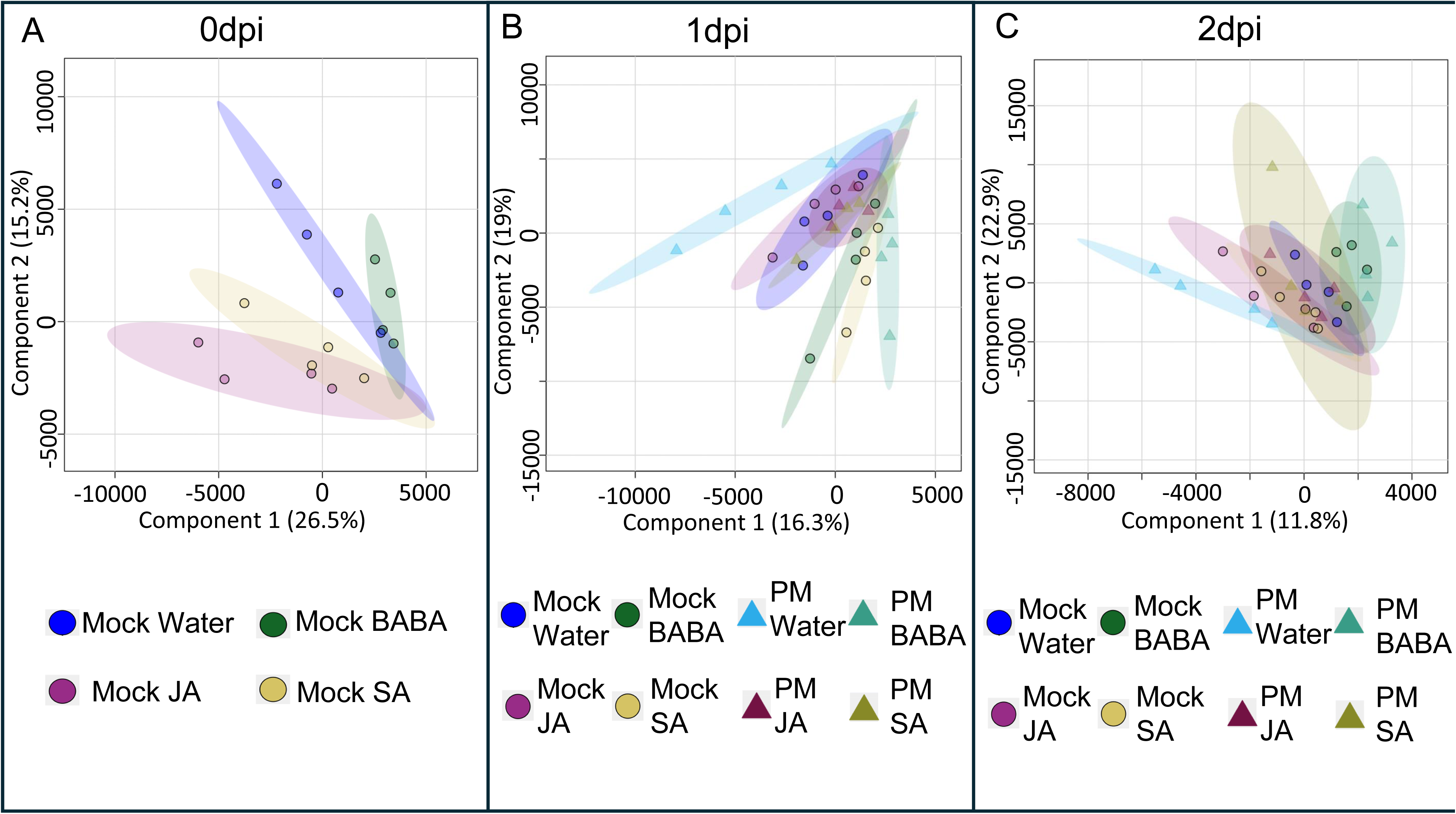
Metabolome PLS-DA plots. A) Partial Least-Squares Discriminant Analysis (PLS-DA) scores plot for mock plants at 0 dpi. B-C) PLS-DA scores plots for mock and infected plants at 1 dpi and 2dpi, respectively.

**Figure S4:**
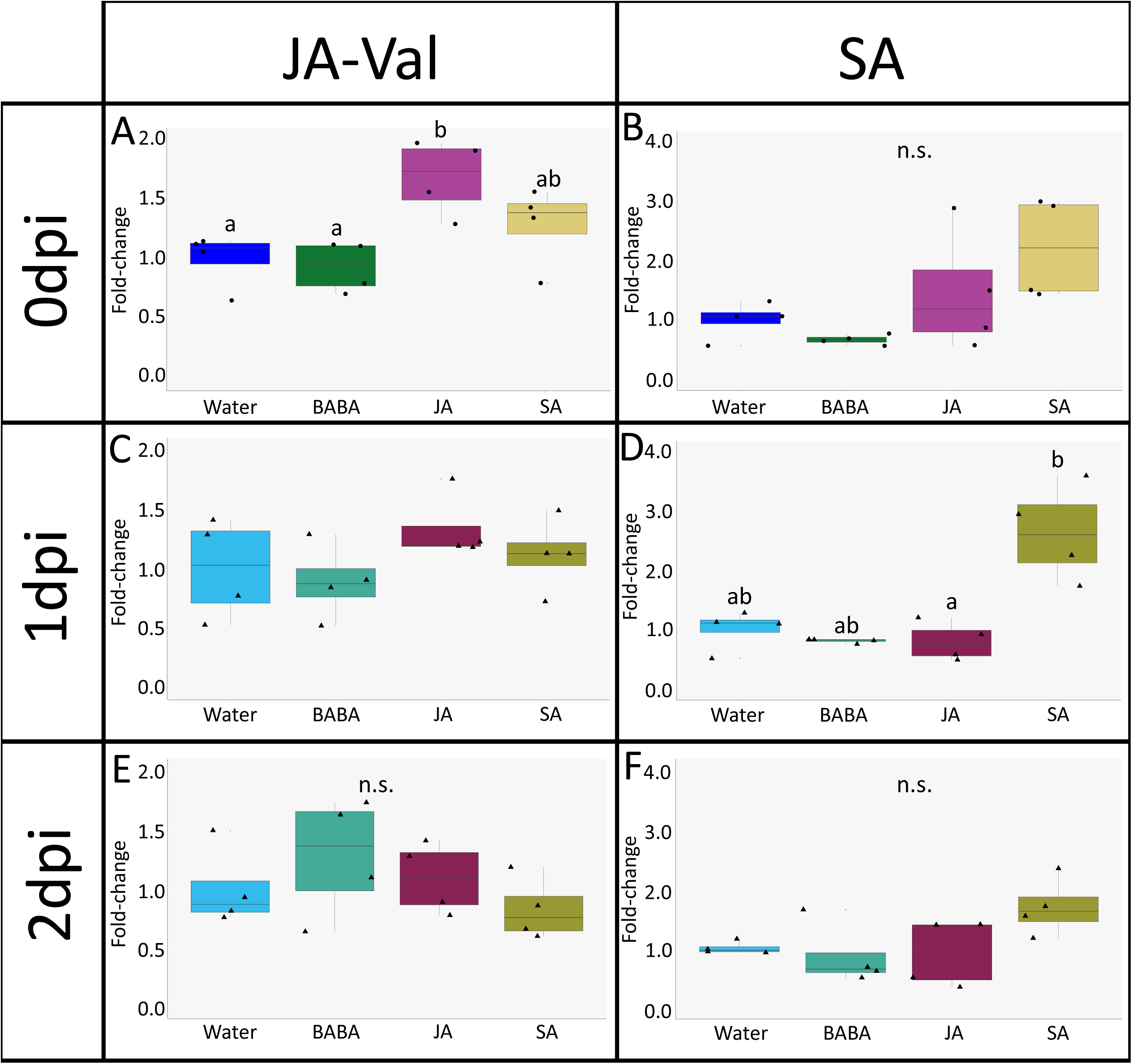
Fold-changes for endogenous hormones following elicitor treatment. A) JA-Val fold-changes compared to water mock at 0 dpi. Letters represent statistically significant differences (ANOVA + Tukey post hoc test, p ≤ 0.05, n = 4). B) SA fold-changes compared to water mock at 0 dpi. Welch ANOVA was not significant (p > 0.05, n = 4). C) JA-Val fold-changes compared to water PM at 1dpi. ANOVA was not significant (p > 0.05, n = 4). D) SA fold-changes compared to water PM at 1dpi. Letters represent statistically significant differences (Welch ANOVA + Dunnet’s T3, p ≤ 0.05, n = 4). E) JA-Val fold-changes compared to water PM at 2dpi. ANOVA was not significant (p > 0.05, n = 4). F) SA fold-changes compared to water PM at 1dpi. ANOVA was not significant (p > 0.05, n = 4). JA-Val = jasmonic acid-valine. SA = salicylic acid.

